# Addressing SARS-CoV-2 Evolution: Neutralization of Emerging Variants of Concern by the AVX/COVID-12 ‘Patria’ Vaccine Based on HexaPro-S Ancestral Wuhan Spike or Its Updated BA.2.75.2 Version

**DOI:** 10.1101/2025.01.23.634579

**Authors:** Gregorio Carballo-Uicab, Gabriela Mellado-Sánchez, Edith González-González, Juana Salinas-Trujano, Ivette Mendoza-Salazar, Karina López-Olvera, Keyla M. Gómez-Castellano, Ma. Isabel Salazar, Jesús M. Torres-Flores, Héctor Elías Chagoya-Cortés, Georgina Paz-De la Rosa, Ignacio Mena, Oscar Rojas-Martínez, Jesús Horacio Lara Puente, Gustavo Javier Peralta-Sánchez, David Sarfati-Mizrahi, Alejandro Torres-Flores, Weina Sun, Florian Krammer, Adolfo García-Sastre, Peter Palese, Constantino López-Macías, Bernardo Lozano-Dubernard, Sonia M. Pérez-Tapia, Juan C. Almagro

**Author notes:** **Correspondence:** (S.M.P.-T.); (J.C.A.), (ext. 62543) (S.M.P.-T.); (J.C.A.). These authors contributed equally to this work (Gregorio Carballo-Uicab, Gabriela Mellado-Sánchez).

## Abstract

Severe acute respiratory syndrome coronavirus 2 (SARS-CoV-2) remains a global health challenge, causing severe morbidity and mortality, particularly in vulnerable groups such as the elderly, immunocompromised individuals, and those with comorbidities. In low- and middle-income countries (LMICs), vaccine access is hindered by high costs and inequitable distribution. To tackle these issues, Mexico developed the AVX/COVID-12 (V-Wu) vaccine, a recombinant Newcastle disease virus (NDV)-based platform expressing a stabilized ancestral Wuhan spike protein (HexaPro-S). Locally manufactured after rigorous testing and regulatory approval, V-Wu aims to enhance self-sufficiency and equity in immunization. This study evaluates an updated vaccine version, AVX/COVID-12 (V-BA), designed to combat Omicron subvariants by expressing the HexaPro-S protein of BA.2.75.2. Both vaccines were administered intramuscularly in K18-hACE2 transgenic and BALB/c mouse models using a prime-boost regimen. Immunogenicity was analyzed by measuring antibodies against Omicron S proteins BA.2.75.2 and XBB.1.5, as well as neutralizing antibodies against Wuhan, BA.1, XBB.1.16, and JN.1 variants. Both vaccines were safe, eliciting robust antibody responses against Omicron S proteins and neutralizing antibodies against multiple emerging SARS-CoV-2 variants of concern (VOCs). V-BA demonstrated superior protection against current Omicron variants, while V-Wu offered broader coverage, including the ancestral Wuhan strain and emerging variants like JN.1. These findings underscore the adaptability of NDV-based platforms in addressing the evolving SARS-CoV-2 landscape and reaffirm the ongoing utility of the ancestral Patria vaccine. Together, they demonstrate the potential of these platforms to drive the development of next-generation vaccines tailored to emerging viral threats, contributing to global health equity.

## Introduction

Since the beginning of the coronavirus 2019 (COVID-19) pandemic, severe acute respiratory syndrome coronavirus 2 (SARS-CoV-2) and its variants have caused millions of deaths and continue to drive significant morbidity (1), particularly in vulnerable populations such as individuals with comorbidities, the immunocompromised, and the elderly. The rapid global spread of the virus has underscored the critical need for effective vaccination campaigns to reduce disease burden and long-term complications. While first-generation vaccines have significantly mitigated the pandemic’s impact, the ongoing emergence of SARS-CoV-2 variants of concern (VOCs) has diminished their effectiveness, making the development of improved vaccines a public health priority (2).

Access to vaccines has posed a significant challenge, particularly in low- and middle-income countries (LMICs), where inequitable distribution and high costs have hindered widespread immunization (3–5). In Mexico, these barriers highlighted the urgent need to develop and produce vaccines locally to achieve self-sufficiency and ensure equitable access for the population. To address this challenge, a multidisciplinary collaboration between the Mexican company AVIMEX, the Icahn School of Medicine at Mount Sinai (USA), Mexican government agencies, and various healthcare and academic institutions resulted in the development of the AVX/COVID-12 vaccine, known as “Patria” (6–8).

The AVX/COVID-12 vaccine is based on the recombinant NDV-platform and expresses the stabilized prefusion S protein (HexaPro-S) of the SARS-CoV-2 ancestral Wuhan strain (9). Preclinical studies demonstrated its safety and immunogenicity, inducing specific antibody and cellular immune responses, including cross-reactivity with some VOCs (10, 11). Phase I, II, and II/III clinical trials (6–8) further validated its safety and immunogenicity when administered via intranasal or intramuscular routes. These findings positioned AVX/COVID-12 as a promising booster vaccine against the original SARS-CoV-2 strain and resulted in regulatory approval for its use in adults in Mexico (12).

However, the ongoing evolution of SARS-CoV-2 underscores the need to update existing vaccines to preserve their efficacy against emerging VOCs. In this study, we evaluated the original AVX/COVID-12 vaccine alongside a newly developed version designed to express the S protein of the Omicron BA.2.75.2 sublineage. The findings and their implications for vaccination strategies using AVX/COVID-12 and its updated formulation to protect against emerging VOCs are discussed.

## Materials and Methods

### Vaccine candidates

The vaccines evaluated in this study were based on a live recombinant NDV vector expressing the S protein of SARS-CoV-2, as previously described in the literature (8, 11, 13). The AVX/COVID-12 vaccine (V-Wu) expresses the S protein of the ancestral Wuhan-1 strain, while the updated NDV-platform vaccine (V-BA) displays the S protein of the SARS-CoV-2 Omicron BA.2.75.2 sublineage. Both versions of the S protein incorporate six proline mutations and a deletion of the polybasic cleavage site, stabilizing the S protein in its prefusion conformation. Additionally, the ectodomain of the S protein is fused to the transmembrane and cytoplasmic domains of the NDV fusion protein to ensure optimal incorporation into the NDV particle. The vaccines were produced by Laboratorio Avi-Mex, S.A. de C.V. in Mexico City using embryonated eggs, under good manufacturing practices (GMP) as previously described (8, 11, 13).

### Animal Models and Experimental Design

This study evaluated the safety and immunogenicity of two experimental vaccines using K18-human angiotensin converting enzyme 2 (hACE2) hACE2 transgenic mice and BALB/c mice as animal models, as illustrated in Figure 1. Transgenic mice were purchased from The Jackson Laboratories (Bar Harbor, ME, USA) and BALB/c mice were produced in the bioterium of the Unidad de Desarrollo e Investigación en Bioterapéuticos (UDIBI).

**Figure 1.**
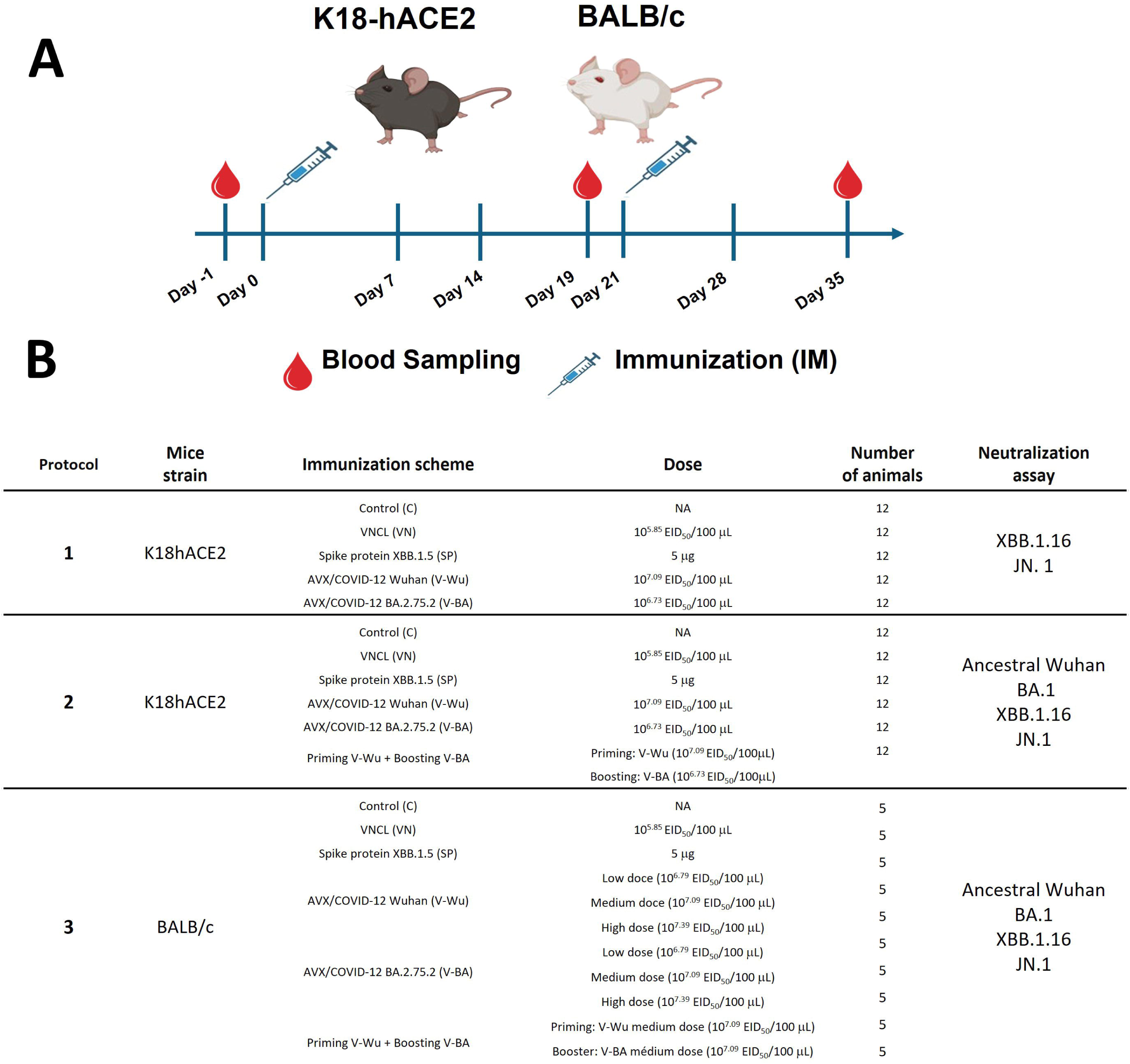
Schematic experimental design. Two mouse models were immunized intramuscularly (IM) with the experimental vaccines AVX/COVID-12 Wuhan (V-Wu) and BA.2.75.2 (V-BA). A) Mice were vaccinated on days 0 and 21. Blood samples were collected on days −1, 19, and 35 via submandibular puncture. B) The table illustrates the different groups divided into three experimental protocols. Protocols 1 and 2 utilized K18-hACE2 mice with a homologous prime-boost immunization scheme, except for group 6 in protocol 2, which received a heterologous prime-boost with V-Wu on day 0 and V-BA on day 21. Protocol 3 analyzed immunization schemes in BALB/c mice, including additional modifications with low (L), medium (M), and high (H) doses of V-Wu and V-BA in a homologous scheme.

The mice were immunized intramuscularly on day 0, followed by a booster dose on day 21. Blood samples were collected at three time points: baseline titers were measured on day −1 (D-1), pre-booster samples were obtained on day 19 (D19), and post-booster samples were collected two weeks later, on day 35 (D35).

Safety: The variation in the body weight of the animals throughout the studies was used as a parameter to monitor mice’s health conditions. It is reported as a percentage relative to the weight at day zero for each animal. Additionally, the skin condition, hair loss, nose moisture, behavioral changes, and posture were monitored three times per week.

All animal care procedures adhered to national regulations (NOM-062-ZOO-1999) and the guidelines of the Office of Laboratory Animal Welfare (OLAW) of the National Institutes of Health (NIH). The animals were housed in standard cages under controlled conditions of temperature and humidity, with a 12-hour light/dark cycle.

### Isolation and identification of SARS-CoV-2 Omicron sublineages

SARS-CoV-2 samples were collected during winter of 2023-2024 from subjects presenting typical COVID-19 symptoms and confirmed by reverse transcription-polymerase chain reaction (RT-PCR), as described by González-González et al. (14). Nasopharyngeal swabs with lowest cycle threshold (Ct) values (<25) were stored in Eagle’s minimum essential medium (EMEM; Manassas, VA, USA; Cat. No. 30-2003). Sample collection adhered to the Declaration of Helsinki principles (15), with informed consent, and all virus handling was conducted under BSL-2+ conditions, following WHO and US Centers of Disease Control and Prevention (CDC) biosafety guidelines (16–18).

Virus isolation: Virus propagation was conducted using Vero cells (ATCC, Cat. No. CCL-81). The cells were incubated with the virus for 60 hours, followed by a 24-hour freezing period to induce cell lysis. Cellular debris was removed by centrifugation, and the supernatants were collected and titrated using a plaque assay (19). RNA was isolated from the culture supernatants with the highest viral titers using the MagMAX Viral and Pathogen Nucleic Acid Isolation Kit (Applied Biosystems™, Thermo Fisher Scientific, Austin, TX, USA; Cat. No. A42352), following the manufacturer’s instructions.

Sanger sequencing of SARS-CoV-2 S1 gene segment: Primary screening was performed to identify the SARS-CoV-2 Omicron XBB.1.16.15 and JN.1 sublineages. cDNA was synthesized from RNA samples using the ProtoScript II First Strand cDNA Synthesis Kit (New England Biolabs, Ipswich, MA, USA). The fragment encoding the S1 protein, spanning nucleotides 950 to 1945 (amino acids 325 to 642), was amplified using the forward primer 5′-ACTTTAGAGTCCAACCAACAGAA-3′ and the reverse primer 5′-AGCCTGCACGTGTTTGAAAA-3′. The PCR reaction performed with Phusion Hot Start Flex DNA Polymerase (New England Biolabs, USA), and size and quality of the amplicons were confirmed by 1% agarose gel electrophoresis. The PCR products were purified from the gel using the QIAquick PCR Purification Kit (Qiagen, Germantown, MD, USA) and subsequently submitted for Sanger sequencing at Wyzer Biosciences, Inc. (Cambridge, MA, USA).

SARS-CoV-2 whole genome sequencing: RNA samples containing the SARS-CoV-2 Omicron XBB.1.16.15 and JN.1 sublineage S1 mutations were submitted for whole-genome sequencing at the Massive Sequencing and Bioinformatics University Unit (UUSMB) of the National Autonomous University of Mexico (UNAM). Sequencing was performed using the COVID-19 Sequencing ARTIC v.5.3.2 Illumina library preparation and the V.5 sequencing protocol (20). The NextSeq500 platform (Illumina, San Diego, CA, USA) was employed with a paired-end read configuration (2 × 150 bp).

The resulting sequences were genotyped using the Pangolin web server (21, 22). All genome sequences achieved a coverage ≥99% and a mean depth ≥1000X. The sequences were deposited in GenBank under the following accession numbers: PP837785.1: AJ153 isolate corresponding to Omicron XBB.1.16.15; PP837746.1: AJ221 isolate corresponding to Omicron BA.1.86; JN.1.

Additionally, this study included the SARS-CoV-2 Wuhan-1 ancestral strain and Omicron BA.1 subvariant, both previously isolated in our laboratory and reported by González-González et al. (14). The genome sequences for these isolates were deposited in GenBank with the following accession numbers: OL790194: SARS-CoV-2 WT and ON651664: SARS-CoV-2 Omicron BA.1.

### Enzyme-linked immunosorbent assay (ELISA) to determine specific IgG against the S protein

NUNC MaxiSorp 96-well flat-bottom ELISA plates (Thermo Scientific, Rochester,NY, USA, Cat. No. 456537) were coated with SARS-CoV-2 XBB.1.5 S Trimer Protein His Tagged (Acro Biosystems, Basel, Switzerland, Cat. No. SPN-C524i) (the same recombinant protein was used as an immunogen admixed 1:1 (v/v) with incomplete Freund’s adjuvant (IFA; Sigma-Aldrich)/100 µL; SP group) or SARS-CoV-2 BA.2.75.2 S Trimer Protein, His Tagged (Acro Biosystems, Cat. No. SPN-C522r) at 2 µg/mL in coating buffer (BioRad, Berkeley, CA, USA, Cat. No. BUF030C) overnight at 2–8 °C. The plates were washed with phosphate buffered saline and 0.1% tween 20 (PBS-T) (PBS 10X, Gibco, Grand Island, NY, USA, Cat. No. 70011-044; Tween 20 Sigma, Darmstadt, Germany, Cat. No. SLCG3047) and blocked with 3% skim milk in PBS-T 0.1% and incubated for 1 hour at room temperature. Subsequently, inactivated serum samples and controls, diluted in 1% milk-PBST, were plated and incubated for 1 hour at room-temperature. The plates were washed with PBS-T and the IgG were detected with horseradish peroxidase (HRP)-conjugated anti-mouse IgG antibody (Invitrogen, Camarillo, CA, USA, Cat. No. 61-6520). The plates were developed by adding the 3,3’,5,5′-tetramethylbenzidine (TMB) substrate (BD OptEIA, San Diego, CA, USA, Cat. No. 555214), and the color reaction was stopped by the addition of TMB stop solution (ABCAM, Waltham, MA, USA, Cat. No. ab171529), and the optical density was measured at 450 nm, with correction at 570 nm, using a plate reader (SpectraMax M3 microplate reader; Molecular Devices, San Jose, CA, USA).

End-titers were reported as ELISA units (EU/mL), the quantification range with linear behavior was established from 0.2–1.4 optical density (OD) values at 450 nm/570 nm using a standard curve generated with a positive serum. The reciprocal of the dilution of the serum, which had a value of 0.2 by interpolation, was established as the lower limit of quantification and was considered as the cutoff point for a positive response.

### Microneutralization assay

Neutralizing antibodies elicited against SARS-CoV-2 were assessed by modifications of the microneutralization method described by Amanat et. al. (23, 24) and the viability method reported by Feoktistova et. al. (24) as follows. Six duplicate dilutions of heat-inactivated serum were prepared in a 96-wells plate, these dilutions were mixed with a constant amount of virus (ancestral isolate or the corresponding Omicron sublineage). Serum-virus complexes were incubated for 1 h at 37°C/5% CO2, then 20,000 Vero cells were placed per well and incubated under the same conditions for 96 h (4 days). After the incubation time of the cells with the serum-virus complexes, culture medium was removed and 10 µL/well of 10% formaldehyde (Sigma Aldrich, St. Louis, MO, USA, Cat. No. 1003643886) in PBS was added and incubated at 4 ° C overnight; the plates were stained with a 0.5% crystal violet solution (Sigma Aldrich) and washed 4 times with running water to remove excess dye; the dye was extracted from the cell monolayer by adding 200 µL/well of methanol (J.T. Baker, USA, Cat. No. 9093-03) and incubating for 20 min at room-temperature (24). After this time, the plates were read in a spectrophotometer at a wavelength of 570 nm. From the neutralization dose-response curves, the 50% inhibitory dilution (ID50) was calculated by applying non-linear regression analysis with GraphPad software.

## Results

### Body weight monitoring and safety evaluation in immunized mice

To evaluate the safety of vaccines V-Wu and V-BA, we monitored weight variations in all experimental groups throughout the study. In K18-hACE2 mice, no significant differences in weight gain were observed, with trends comparable to those of the control group (Figure 2A and B). Similarly, BALB/c mice showed no significant changes in weight gain (Figure 2C). When analyzing intra-group trends, BALB/c mice exhibited slower weight gain compared to K18-hACE2 mice. This difference can be attributed to variations in strain, age, and initial weight. At the start of the study, K18-hACE2 mice weighed 16–18 g and were six weeks old, while BALB/c mice weighed 19–22 g and were seven weeks old. Despite these initial differences, neither vaccine affected normal growth or weight gain in either strain, as demonstrated by comparisons with the unvaccinated control group. Additional clinical observations, including assessments of skin condition, hair quality, nasal moisture, and behavior, revealed no abnormalities in any group during the study.

**Figure 2.**
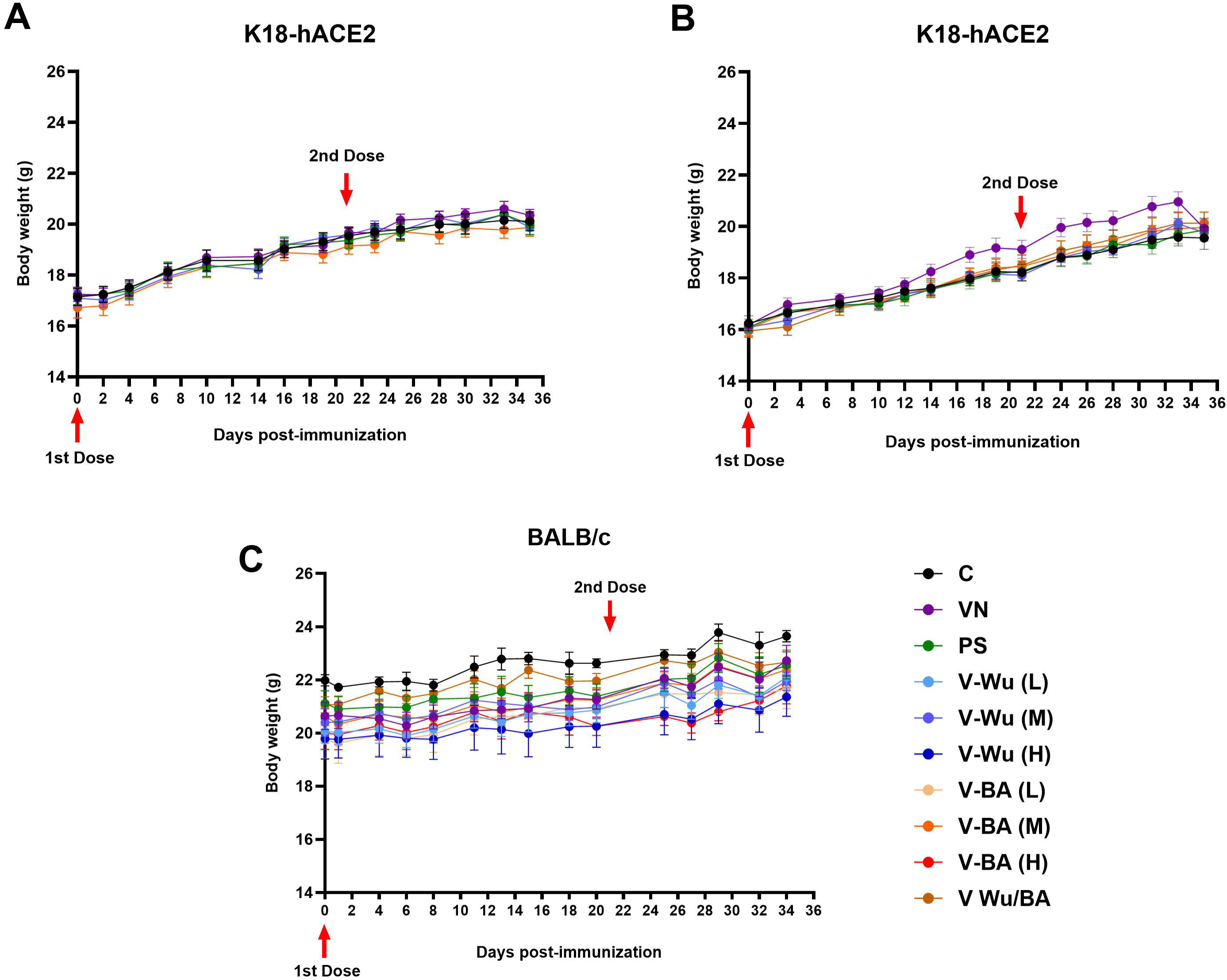
Safety and tolerability of V-Wu and V-BA vaccines post-immunization. Kinetics of body weight in mice following immunization across different experimental protocols. A) K18-hACE2 mice immunized with homologous schemes. B) K18-hACE2 mice immunized with homologous or heterologous schemes (prime with V-Wu and boost with V-BA). C) BALB/c mice immunized with homologous schemes, including variations with low (10^6.79^ EID50/100 µL), medium (10^7.09^ EID50/100 µL), and high (10^7.39^ EID50/100 µL) doses of V-Wu or V-BA, as well as a heterologous scheme (prime with V-Wu at medium dose and boost with V-BA at medium dose). The graphs depict the mean ± SEM of body weight (g) for each study protocol. Red arrows indicate vaccination days for priming (Day 0) and boosting (Day 21). C = Negative control, VN = Vector control, SP = S protein (XBB.1.5 sublineage), V-Wu = AVX/COVID-12 Wuhan vaccine, V-BA = AVX/COVID-12 BA.2.75.2 vaccine, L = Low dose, M = Medium dose, H = High dose, V-Wu/BA = Prime with V-Wu and boost with V-BA. K18-hACE2 groups: n=12 per group; BALB/c groups: n=5 per group. Data were analyzed with two-way ANOVA and Tukey’s post hoc test using GraphPad Prism software. Statistical differences are indicated where p < 0.05.

### V-Wu and V-BA immunization induced specific antibodies against Omicron BA.2.75.2 and XBB.1.5 sublineages

The evaluation of antibody titers induced by homologous immunization with V-Wu or V-BA against the S protein of the Omicron sublineages BA.2.75.2 and XBB.1.5 in K18-hACE2 mice showed no significant differences in antibody titers after priming with V-Wu (Figure 3A and B). However, by day 35, a homologous booster dose with either V-Wu or V-BA resulted in an increase in antibody titers compared to the controls or the empty vector groups (Figure 3A and B). In the BALB/c mouse model, the dependence of the primary response to the administered dose (low, medium, or high) of both vaccines was evaluated. The results revealed antibody titers comparable to those observed in K18-hACE2 mice at both day 19 and day 35, with no significant differences between doses.

**Figure 3.**
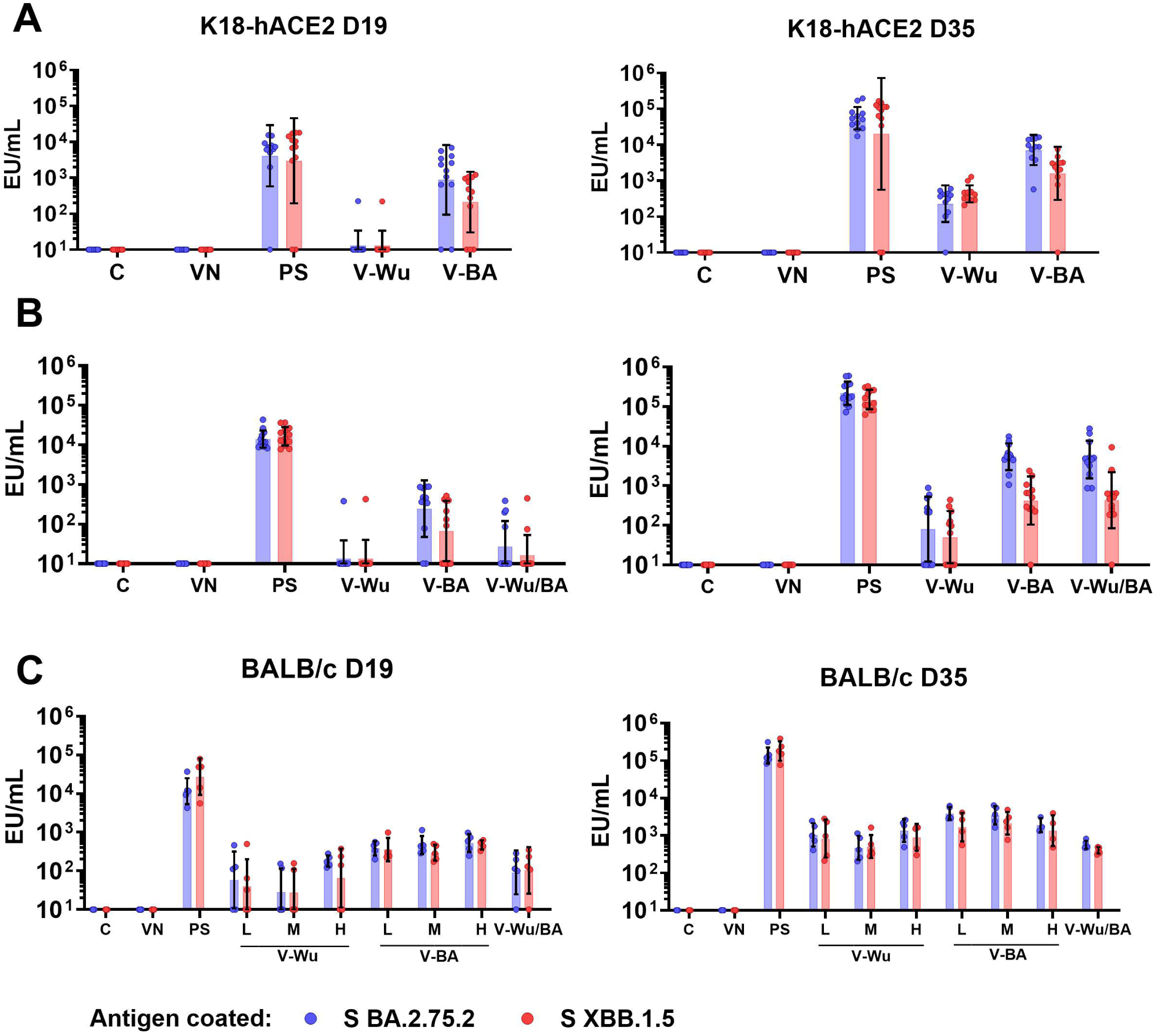
Immunization with V-Wu and V-BA induces specific antibodies against BA.2.75.2 and XBB.1.5 S proteins. Specific IgG antibodies expressed as ELISA units (EU) per mL against the S proteins of SARS-CoV-2 BA.2.75.2 and XBB.1.5 variants were measured by ELISA on days 19 and 35 post-immunization. A) K18-hACE2 mice immunized with homologous schemes. B) K18-hACE2 mice immunized with homologous and heterologous schemes (priming with V-Wu and boosting with V-BA). C) BALB/c mice immunized with homologous schemes, including variations in dose (low: 10^6.79^ EID50/100 µL, medium: 10^7.09^ EID50/100 µL, and high: 10^7.39^ EID50/100 µL) for V-Wu or V-BA, as well as a heterologous scheme (priming with V-Wu at medium dose and boosting with V-BA at medium dose). The graphs display the geometric mean ± SD of EU/mL for each group. C = Negative control, VN = Vector control, SP = S protein (XBB.1.5 sublineage), V-Wu = AVX/COVID-12 Wuhan vaccine, V-BA = AVX/COVID-12 BA.2.75.2 vaccine, L = Low dose, M = Medium dose, H = High dose, V-Wu/BA = Prime with V-Wu and boost with V-BA. Sample sizes: K18-hACE2 groups (n = 12 per group); BALB/c groups (n = 5 per group).

When comparing the heterologous regimen of V-Wu priming followed by V-BA boosting with the homologous V-BA regimen, comparable antibody titers were observed in K18-hACE2 mice (Figure 3B). However, in BALB/c mice, the antibody titers generated by heterologous vaccination were lower and remained comparable between the priming and booster doses (Figure 3C). Overall, both homologous and heterologous vaccination regimens with either vaccine induced comparable antibody titers against the Omicron sublineages BA.2.75.2 and XBB.1.5.

### V-Wu and V-BA induced neutralizing antibody responses against the ancestral strain and Omicron sublineages

Similar antibody titers were induced by the two vaccines, suggesting comparable overall immune activation. Despite this, the two vaccines exhibited differences in neutralizing activity against the Omicron sublineages. Neutralizing antibody titers were evaluated on day 35 following homologous V-Wu and V-BA vaccination schemes, as shown in dilution curves and ID50 values for K18-hACE2 mice groups (Figure 4). V-Wu did not elicit neutralizing antibodies against the XBB.1.16 strain. However, it elicited a statistically significant response against the JN.1 variant compared to the control groups (Figure 4A).

**Figure 4.**
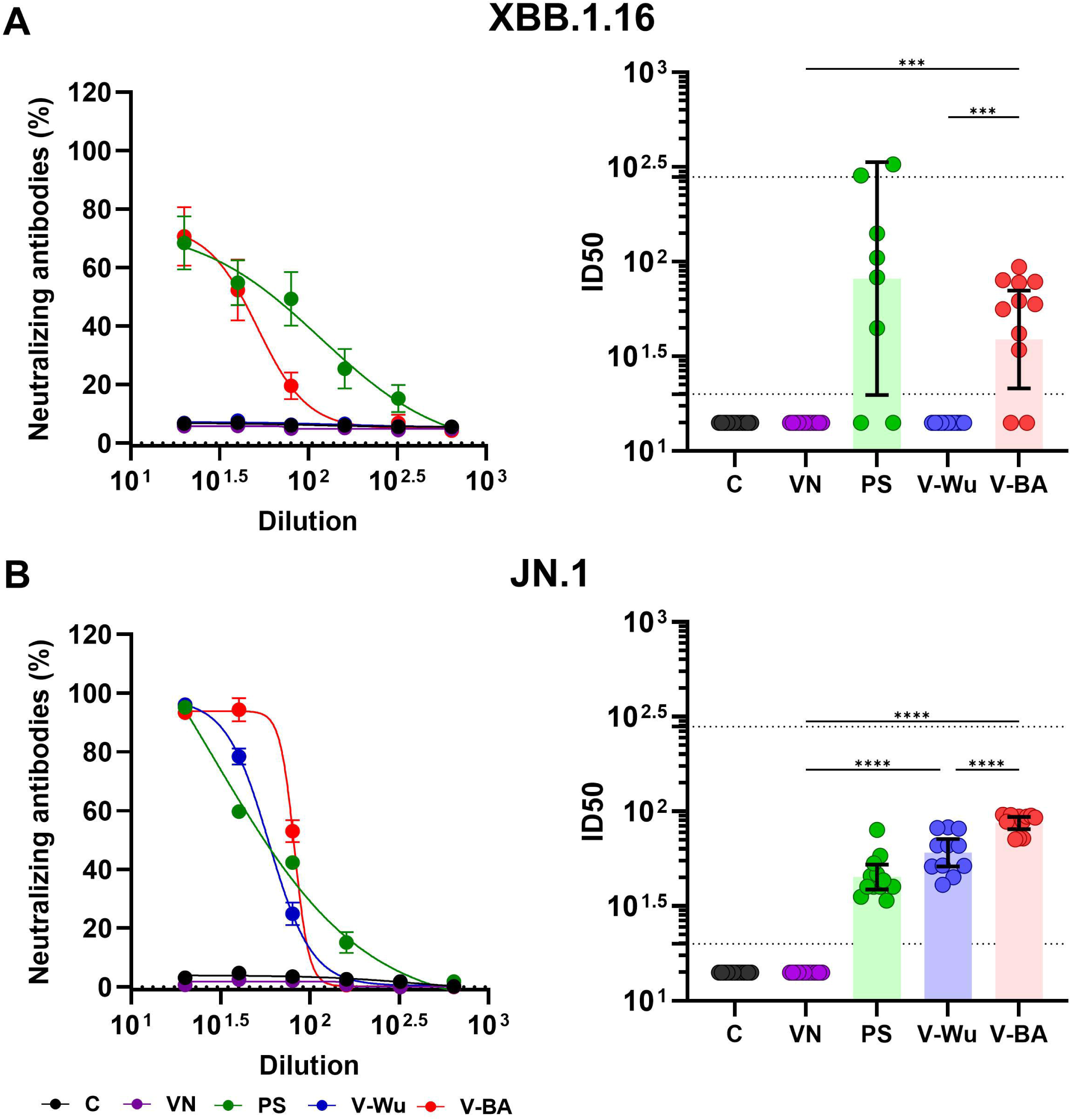
Neutralizing antibody responses against XBB.1.16 and JN.1 Omicron variants induced by homologous immunization with V-Wu or V-BA vaccines. Neutralizing activity of sera from K18-hACE2 mice immunized with homologous vaccine schemes was assessed against SARS-CoV-2 variants A) XBB.1.16 or B) JN.1 Omicron sublineages. The left panels display the neutralization percentage (mean ± SEM) derived from serial dilutions of sera collected on day 35 post-boost. The right panels show the ID50 values (mean ± 95% CI) calculated at the same time point. Dotted lines indicate the assay’s detection limit (lower line) and the response level of the positive control (upper line). Groups: C = Negative control, VN = Vector control, SP = S protein (XBB.1.5 sublineage), V-Wu = AVX/COVID-12 Wuhan vaccine, V-BA = AVX/COVID-12 BA.2.75.2 vaccine. Sample size: n = 12 per group. Data were analyzed with one-way ANOVA and Tukey’s post hoc test using GraphPad Prism software. Statistical significance is denoted where ****p < 0.0001.

Immunization with V-BA resulted in a statistically significant increase in neutralizing antibodies against both Omicron sublineages XBB.1.16 (Figure 4B) and JN.1, compared to V-Wu. The control groups and the vector did not elicit neutralizing antibodies. In contrast, recombinant XBB.1.5 S protein immunization showed a response for both Omicron sublineages.

We also compared homologous and heterologous immunization (Figure 5). V-Wu elicited statistically significant increases in neutralizing antibodies against the ancestral strain compared to V-BA (Fig. 5A), consistent with previous analyses (Figure 4). V-Wu also induced neutralizing antibodies against JN.1 but did not generate a neutralizing antibody response against XBB.1.16 (Fig. 5C) and exhibited a weak and variable response against BA.1 (Fig. 5B and D). On the other hand, V-BA elicited neutralizing antibodies against Omicron BA.1, XBB.1.16, and JN.1 sublineages, but not against the ancestral lineage (Figure 5). The heterologous vaccination induced neutralizing antibodies against ancestral and the Omicron sublineages BA.1, XBB.1.16, and JN.1.

**Figure 5.**
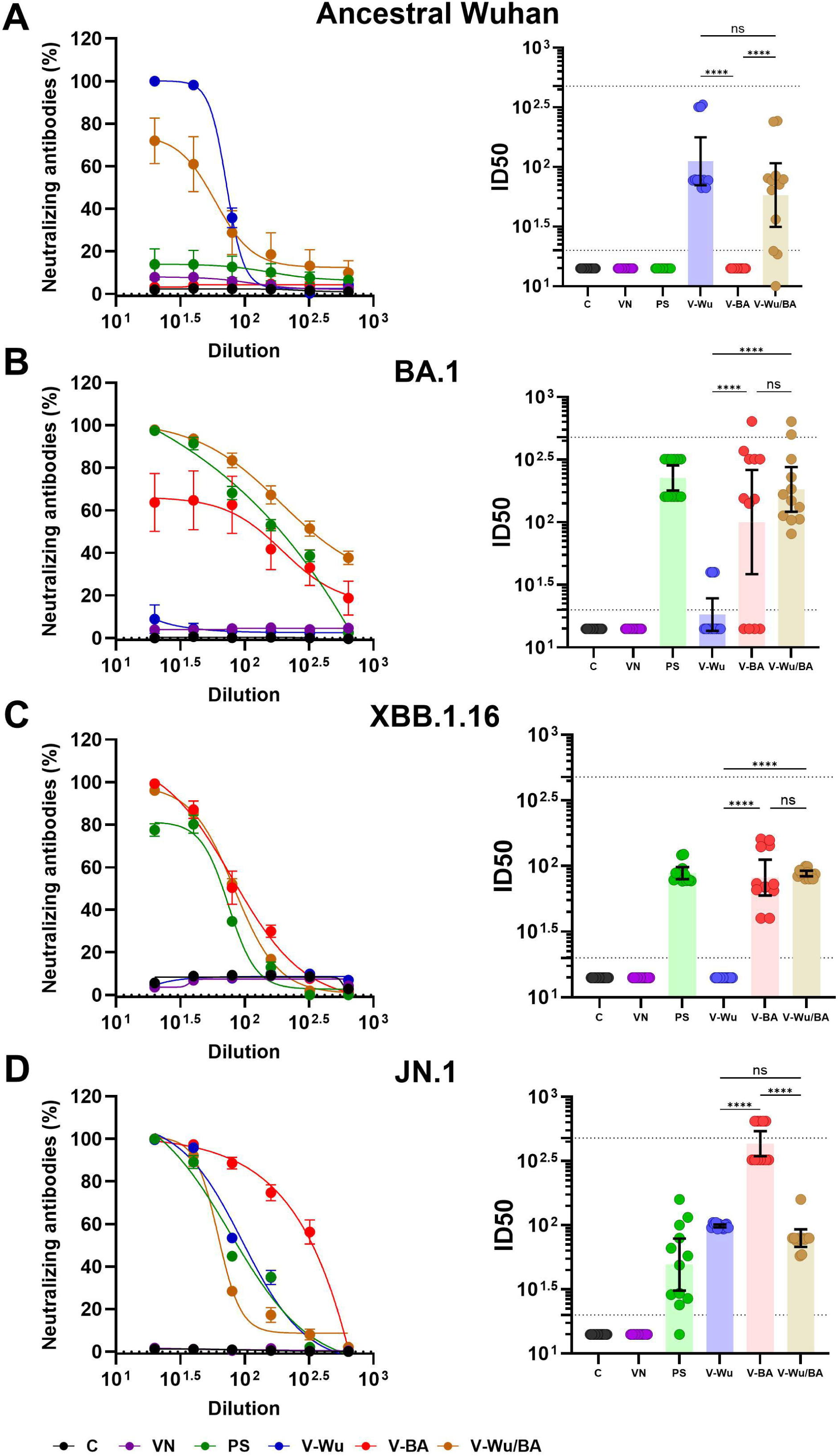
Neutralizing antibody responses against ancestral Wuhan and Omicron variants induced by homologous and heterologous immunization with V-Wu or V-BA vaccines. Neutralizing activity of sera from K18-hACE2 mice immunized with homologous (V-Wu/Wu or V-BA/BA) or heterologous (V-Wu/BA) vaccine schemes was evaluated against SARS-CoV-2 variants: A) ancestral Wuhan-1, B) BA.1, C) XBB.1.16, and D) JN.1 Omicron sublineages. The left panels illustrate neutralization percentages (mean ± SEM) obtained from serial dilutions of sera collected on day 35 post-boost. The right panels present ID50 values (mean ± 95% CI) calculated for the same time point. Dotted lines represent the assay’s detection limit (lower line) and the response level of the positive control (upper line). Groups: C = Negative control, VN = Vector control, SP = S protein (XBB.1.5 sublineage), V-Wu = AVX/COVID-12 Wuhan vaccine, V-BA = AVX/COVID-12 BA.2.75.2 vaccine, V-Wu/BA = Prime with V-Wu and boost with V-BA. Sample size: n = 12 per group. Data were analyzed with one-way ANOVA and Tukey’s post hoc test using GraphPad Prism software. Statistical significance is denoted as ****p < 0.0001, ***p < 0.001, and ns = not significant.

When analyzing the neutralizing response in BALB/c mice, the results were consistent with those showed for the K18-ACE2 groups (Figure 6). V-Wu exhibited a dose-dependent increase in neutralizing antibodies against both ancestral and Omicron JN.1 strains (Figure 6A and D). It also elicited neutralizing antibodies against BA.1 sublineage but not against XBB.1.16 (Figure 6B and C). Interestingly, the high-dose V-Wu group developed a neutralizing response similar to that observed in the low and medium-dose V-BA groups. On the other hand, the heterologous vaccination induced neutralizing antibodies against all SARS-CoV-2 strains (Figure 6), further supporting the use of the heterologous scheme to cover the variants analyzed in this study.

**Figure 6.**
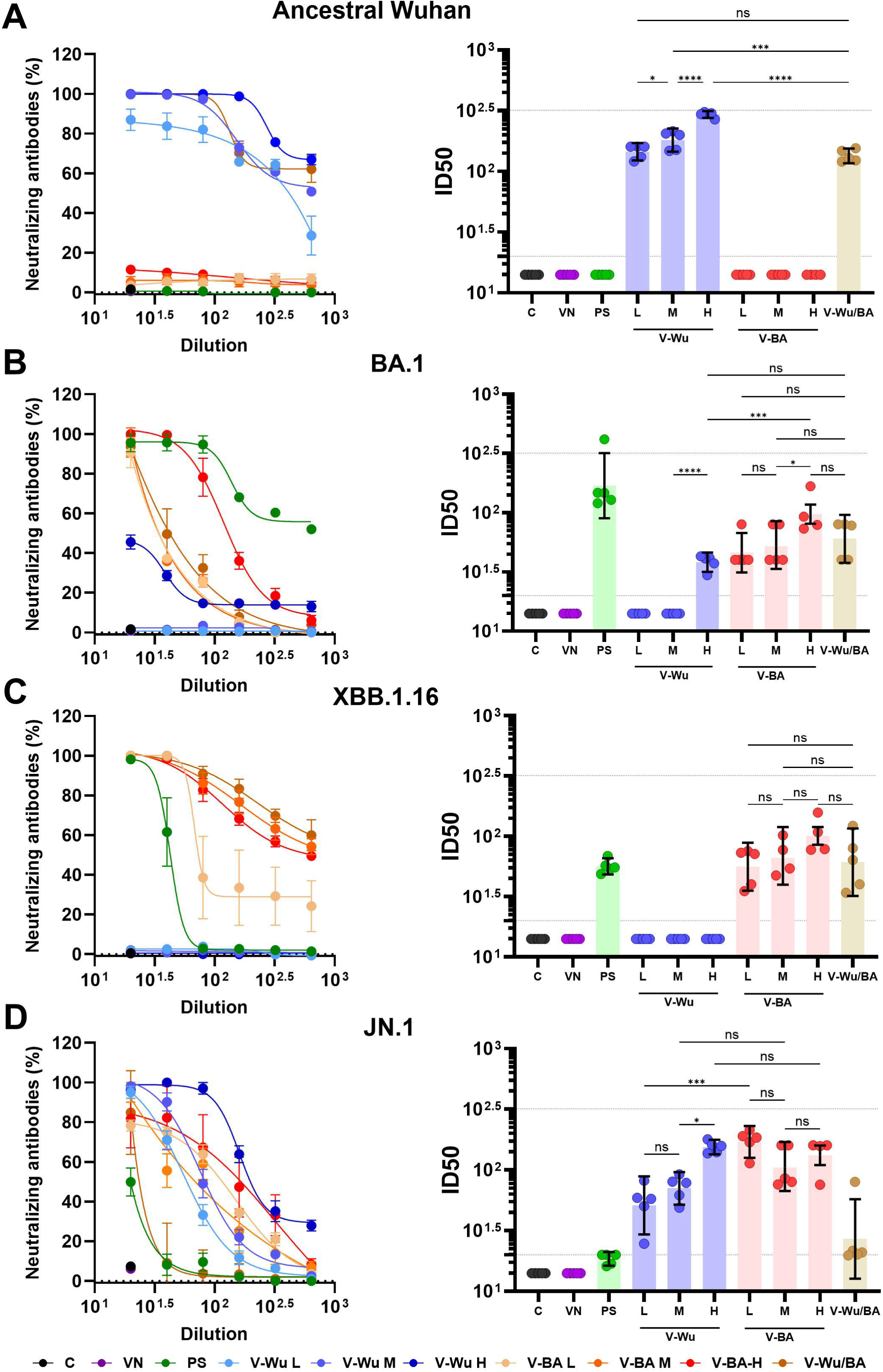
Neutralizing antibody responses against ancestral Wuhan and Omicron variants induced by dose-response homologous or heterologous immunization with V-Wu or V-BA vaccines. Neutralizing activity of sera from BALB/c mice immunized with homologous (V-Wu/Wu or V-BA/BA) schemes, including variations in dose (low: 10^6.79^ EID50/100 µL, medium: 10^7.09^ EID50/100 µL, and high: 10^7.39^ EID50/100 µL), or heterologous (V-Wu/BA) vaccine schemes was assessed against SARS-CoV-2 variants: A) ancestral Wuhan-1, B) BA.1, C) XBB.1.16, and D) JN.1 Omicron sublineages. The left panels display neutralization percentages (mean ± SEM) derived from serial dilutions of sera collected on day 35 post-boost, while the right panels present ID50 values (mean ± 95% CI) calculated for the same time point. Dotted lines indicate the assay’s detection limit (lower line) and the response level of the positive control (upper line). C = Negative control, VN = Vector control, SP = S protein (XBB.1.5 variant), V-Wu = AVX/COVID-12 Wuhan vaccine, V-BA = AVX/COVID-12 BA.2.75.2 vaccine, L = Low dose, M = Medium dose, H = High dose, V-Wu/BA = Prime with V-Wu and boost with V-BA. Sample size: n = 5 per group. Data were analyzed with one-way ANOVA and Tukey’s post hoc test using GraphPad Prism software. Statistical significance is denoted as ****p < 0.0001, ***p < 0.001, and ns = not significant.

## Discussion

The V-Wu vaccine, based on the ancestral Wuhan strain, has successfully demonstrated safety, tolerability, and immunogenicity in phase 1-3 clinical trials (6–8). Additionally, a non-inferiority study revealed that the antibody responses induced by V-Wu are comparable to those elicited by the ChAdOx-1-S vaccine from AstraZeneca (6). As a result, V-Wu received regulatory approval for use as a booster in Mexico (12). However, the rapid evolution of SARS-CoV-2 has led vaccine manufacturers to update formulations to address prevalent variants. Unfortunately, the pace of viral evolution outstrips our ability to develop and test vaccines tailored to the currently VOCs, thus underscoring the importance of creating vaccine formulations capable of eliciting a broader immune response to combat multiple variants effectively.

In this context, the development of the V-BA vaccine aimed to target broader conserved epitopes shared among various circulating strains. BA.2.75.2 was selected as the basis for this formulation, as it serves as a close common ancestor for most of the currently circulating variants (19). The results of the preclinical trial in mice described in this report demonstrated that the V-BA vaccine is safe and well-tolerated, comparable to V-Wu. Both K18-hACE2 and BALB/c mice exhibited favorable responses without evident adverse effects on overall health.

In Mexico, most available vaccines are derived from the ancestral Wuhan strain, and the majority of the population has been vaccinated exclusively with these, including V-Wu. Consequently, we evaluated the immune response induced by priming with V-Wu and boosting with V-BA as an experimental model to simulate the potential application of V-BA in a population previously vaccinated with first-generation COVID-19 vaccines. Our results confirmed that this combination was also safe and well-tolerated.

Furthermore, the V-BA vaccine induced robust antibody responses in both K18-hACE2 and BALB/c mice, comparable to those elicited by V-Wu, demonstrating that V-BA is highly immunogenic. Interestingly, V-BA elicited neutralizing antibodies against the Omicron BA.2.75.2, XBB.1.5, and JN.1 sublineages but failed to generate a neutralizing response against the ancestral SARS-CoV-2 strain. This lack of response may be attributed to the significant genetic divergence of Omicron sublineages from the ancestral strain. The first Omicron sublineage (BA.1), reported in November 2021 (25), represented the most substantial deviation from Wuhan among all SARS-CoV-2 VOCs, introducing a major shift in immune evasion. BA.1 featured five deletions and 34 mutations in the S protein (26), with 15 mutations in the receptor-biding domain (RBD) (lower part of Figure 7), which dramatically altered its antigenic properties. Notably, most prophylactic or therapeutic antibodies previously approved by the Food and Drug Administration for treating COVID-19 lost efficacy when challenged with Omicron BA.1 (27). Therefore, it was not necessarily expected that a vaccine incorporating the S protein from Omicron BA.2.75.2 would induce neutralizing antibodies against the ancestral Wuhan strain.

**Figure 7.**
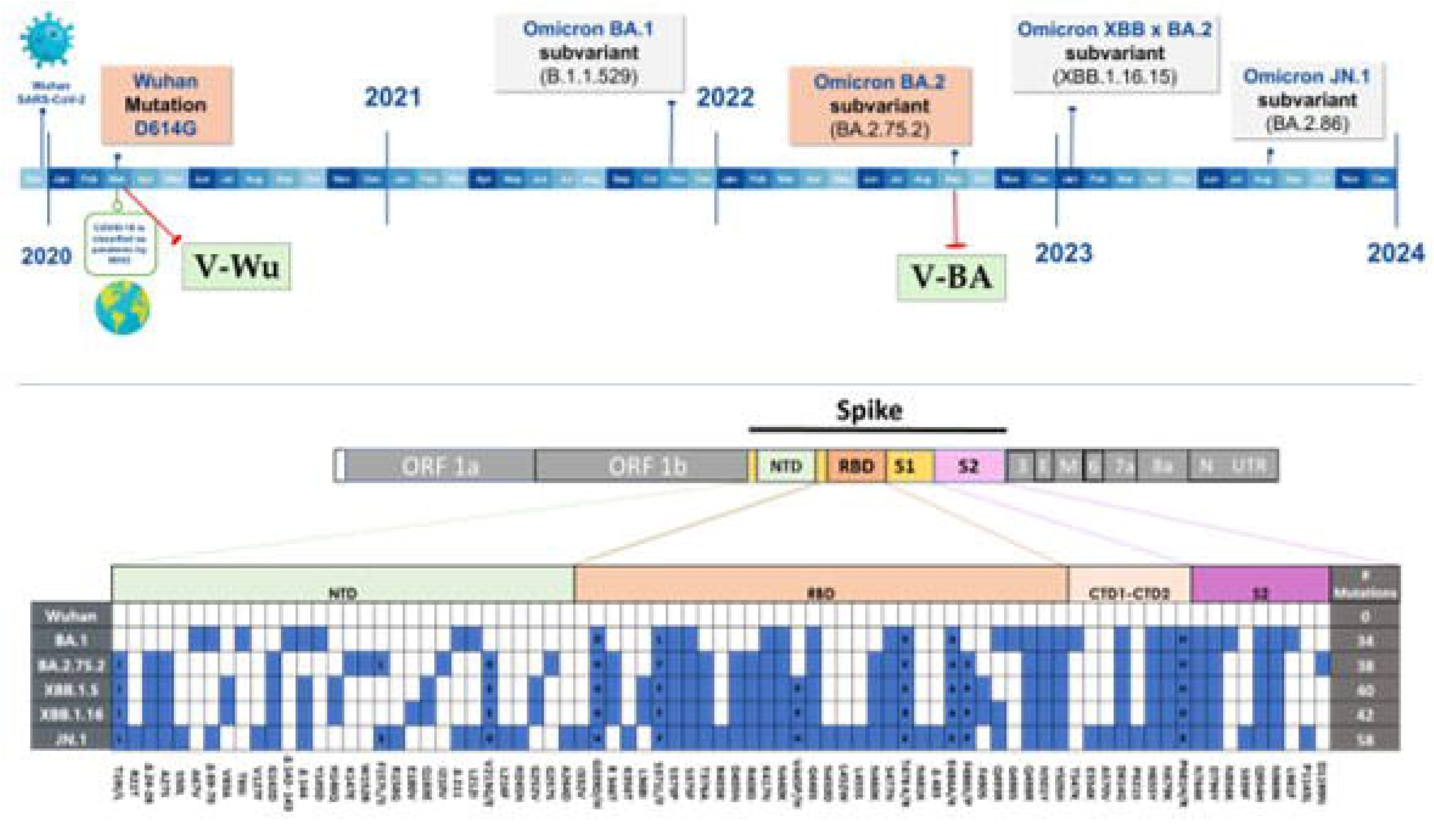
SARS-CoV-2 evolution. Upper panel: Timeline illustrating the emergence of SARS-CoV-2 and Omicron variants. Variants of the S protein incorporated into vaccine designs are highlighted in red boxes, while additional S protein variants used in ELISA or neutralization assays are shown in grey boxes. Lower panel: Mutations (blue squares) in the S protein of Omicron subvariants relative to the ancestral Wuhan-1 virus.

The fact that V-Wu elicited specific antibodies against the S protein of Omicron BA.2.75.2 and XBB.1.5, following both homologous boosting with V-Wu and heterologous boosting with V-BA, could be attributed to their evolutionary relationship with the BA.1 and XBB.1.16 sublineages (25, 26). The ability of V-Wu to generate a strong neutralizing response against JN.1 but only weak titers against BA.1 and no response against XBB.1.16 may be due to several factors, including antigen specificity, revertant mutations in the S protein that made Wuhan-based immunization more effective against JN.1 than BA.1 and XBB.1.16, differences in dosage, immunological variability among individuals, and the sensitivity of the microneutralization assay used to detect neutralizing antibodies against BA.1 and XBB.1.16.

Another explanation for this cross-reactivity could be related to the structural modifications in the S protein which enhance both its stability and immunogenic potential as been reported previously (9, 28). Both vaccines leverage an NDV platform incorporating HexaPro stabilization, which maintains class I fusion proteins in their prefusion conformation (9). This strategy has proven pivotal in exposing a broad range of epitopes in both open and closed structural conformations, thereby enhancing neutralizing responses (9, 28). The efficacy of similar stabilization techniques is evident in recently approved vaccines for respiratory syncytial virus (RSV) (29). Moreover, advanced stabilization methods, such as disulfide bond introduction and cavity-filling substitutions, have been successfully applied to other vaccine candidates, including those developed for RSV, ebolavirus, influenza virus and human immunodeficiency virus 1 (29–32).

Building on these technological advancements, the V-Wu and V-BA vaccines exhibit a robust capacity to trigger broad immunogenic responses, underscoring their potential to combat emerging SARS-CoV-2 variants. Preclinical and clinical trials using V-Wu have demonstrated the induction of high antibody titers and T-cell activation against both ancestral and VOCs following immunization and/or boosting (7, 8, 11). In vitro antigenicity assays with the V-Wu vaccine showed its ability to restimulate cellular responses, where T-cells recognized conserved epitopes encoded by the vaccine in individuals who had recovered from COVID-19 or were vaccinated with other vaccine platforms (13). Additionally, these epitopes exhibited cross-reactivity with antibodies generated in response to the ancestral Wuhan strain and early Omicron variants, highlighting the conserved B- and T-cell epitopes across SARS-CoV-2 variants (13, 33).

Finally, it should be highlighted that our results have significant implications for the development of new vaccines and the design of vaccination strategies. First, priming and boosting with V-Wu appeared to elicit a robust neutralizing response against both the ancestral strain and recent SARS-CoV-2 VOCs, particularly JN.1 and BA.1 (especially with a high dose). Given the rapid evolution of SARS-CoV-2 and the time and resources required to develop and test new vaccines, our findings suggest that V-Wu, already successfully tested in clinical trials, could serve as an effective booster to provide protection against recent VOCs. Second, increasing the dose may further enhance the breadth of the neutralizing response. Third, heterologous vaccination demonstrated the ability to induce neutralizing antibodies against all the SARS-CoV-2 strains tested and thus, priming with V-Wu followed by a V-BA booster may offer additional benefits in enhancing and broadening the antibody responses against emergent VOCs.

In conclusion, the results of this study demonstrate that V-BA is safe, well-tolerated, and immunogenic in mice, inducing robust and broad neutralizing antibody responses against various Omicron sublineages, both as a homologous booster and in combination with V-Wu as a heterologous boost. This makes V-BA a promising candidate to update the V-Wu vaccine for use as a booster in the population. In addition, these findings underscore the adaptability of NDV-based platforms in addressing the evolving SARS-CoV-2 landscape and reaffirm the ongoing utility of the ancestral Patria vaccine. Together, they demonstrate the potential of these platforms to drive the development of next-generation vaccines tailored to emerging viral threats, contributing to global health equity.

## Author Contributions

Conceptualization, design, execution of most of the experimental strategies, analysis, interpretation of the results, and manuscript writing, G.C.-U., G.M.-S., H.E.C.-C., and J.C.-A; performed neutralization assays, validated the experiments, supervised the work, and data analysis, E.G.-G. and J.S.-T.; validated methods, quantified IgG in serum samples and data analysis, I.M.-S.; animal handling, sampling for various analyses and data analysis, K.L.-O.; sequence analysis of the different variants, K.G.-C.; generating the results, preparing the figures, discussing the findings, and reviewing the manuscript, G.C.-U., G.M.-S., E.G.-G., J.S.-T., I.M.-S., K.L.-O., and K.G.-C.; conceptualized the manuscript and contributed to the overall direction of the work, manuscript review, and editing, S.M.P.-T. and J.C.-A.; data curation, writing - review & editing, C.L.-M and A.T.-F.; acquired funding, G. P.-D., O R.-M., J.H.L.-P., G.J.P.-S., D. S.-M., B. L.-D; participated in the review, commentary, and revision of the manuscript, M.I.S., J.M.T.-F., A.G.-S., F.K., W.S. and P.P. All authors have read and agreed to the final version of the manuscript.

## Acknowledgments

Our gratitude goes to Blanca Jazmín Sánchez-Morales and Eduardo Ivan Aguilar-Salgado for the Quality Assurance management, to Luis Alberto Valencia-Flores, Sandra Comparán-Alarcón, Efrén Ernesto Enríquez-Pérez, Jesús Olvera-Flores, Mireya Ramírez Florencio, Guillermo Alejandro Islas Saldívar, Stephany Daniela Rodríguez-Luna, Giovanni Santiago-Casas and the rest of the personnel of In vivo models, Molecular Biology-Discovery, Biological Characterization, Virology, Quality Assurance, and Biobank areas of UDIBI for their technical support. We also would like to thank Dr. Martha Torres-Rojas and Dr. Horacio Zamudio from Instituto Nacional de Enfermedades Respiratorias “Ismael Cosío Villegas” (INER) for providing us with samples to validate the microneutralization test. Finally, we thank Ghersia Contreras-Esparza, Ignacio Mejia-Calvo and Alejandro Ruiz Martínez for their administrative support of the project.

## Institutional Review Board Statement

The study was conducted in accordance with the Declaration of Helsinki and approved by the Research Ethics Committee of the Escuela Nacional de Ciencias Biológicas (CEI-ENCB) (protocol code ZOO-021-2023, approved Dec 21st, 2023).

